# Manatee: detection and quantification of small non-coding RNAs from next-generation sequencing data

**DOI:** 10.1101/662007

**Authors:** Joanna E Handzlik, Spyros Tastsoglou, Ioannis S Vlachos, Artemis G Hatzigeorgiou

## Abstract

Small non-coding RNAs (sncRNAs) play important roles in health and disease. Next Generation Sequencing technologies are considered as the most powerful and versatile methodologies to explore small RNA (sRNA) transcriptomes in diverse experimental and clinical studies. Small RNA-Seq data analysis proved to be challenging due to non-unique genomic origin, short length and abundant post-transcriptional modifications of sRNA species. Here we present Manatee, an algorithm for quantification of sRNA classes and detection of uncharacterized expressed non-coding loci. Manatee adopts a novel approach for abundance estimation of genomic reads that combines sRNA annotation with reliable alignment density information and extensive reads salvation. Comparison of Manatee against state-of-the-art implementations using real/simulated data sets demonstrates its superior accuracy in quantification of diverse sRNA classes providing at the same time insights about unannotated expressed loci. It is user-friendly, easily embeddable in pipelines and provides a simplified output suitable for direct usage in downstream analyses and functional studies.

## Introduction

The discovery of short functional RNA classes such as microRNAs (miRNAs) and small interfering RNAs (siRNAs) revealed their involvement in pervasive regulation of gene expression and inaugurated the RNA revolution. Next Generation Sequencing (NGS) techniques offer a powerful high-throughput means for the quantification and discovery of many small RNA classes (1). Small RNA sequencing (sRNA-Seq) has been established as the gold standard technique for high-throughput detection and quantification of small RNAs typically between 18 and 35 nucleotides in length, enabling expression studies of small RNA species, as well as the discovery of novel small non-coding RNAs. miRNAs have been the focal point of such analyses, since they play a pivotal role in post-transcriptional regulation of gene expression (2) influencing development and disease pathogenesis (3, 4). Other sRNAs identified in NGS experiments, such as ribosomal RNAs (rRNAs), transfer RNAs (tRNAs) and small nucleolar RNAs (snoRNAs), were usually conceived as findings of secondary significance. However, recent studies have provided insight into novel biological roles of such sRNAs (5–7). Using relevant approaches, new short RNA families with biological functions that are still under debate have been discovered. tRNA-derived RNA fragments (tRFs), a novel class of sRNAs second in abundance only to miRNAs (5) or box C/D snoRNAs (7) comprise characteristic examples of such classes. The majority of tRF sequences are derived from precise cleavage and processing at the 5′ or 3′ end of mature or precursor tRNAs, and studies indicate their possible involvement in miRNA-like RNA targeting as well as global translational suppression (8). snoRNAs, known to serve functions in RNA modification processes (9), have been recently shown to host specific miRNA-like short RNAs and have been found deregulated in various diseases and malignancies (6, 7). Hence, accurate quantification and analysis of the full sRNA spectrum is of great interest.

Currently the analysis of sRNA-Seq data is not as mature as for longer RNAs, and their usefulness is impacted by major hindrances. Particularly, the short length (usually ∼18-30nt) of sRNA-Seq reads introduces the problem of multi-mapping, where reads align to more than one genomic locations with equal alignment scores. This issue is exacerbated if we consider that many sRNAs are transcribed from repeat loci (10). As a consequence, the most common approaches adopted for RNA-Seq data (11) cannot be successfully applied to this setting; retaining only uniquely aligned reads (12) leads to the omission of a significant portion of reads, while equal distribution (13, 14), random read placement (15) or reporting all possible alignment positions of multi-mapping reads (16), inevitably leads to incorrect or indirectly quantifiable results (11). Additionally, the existence of numerous intermediate and terminal products of sRNA biogenesis and potentially of yet unknown RNA species in sRNA-Seq data is undermined with current approaches (17).

State-of-the-art methods employ direct alignment against known miRNA or sRNA annotations, in order to diminish the extent of multi-mapping (11). However, these methods are bound to quantifying only known sRNAs, while reads that could align better in other genomic places are forced to map with lower scores in the reduced search space (18). The ambiguity of the genomic origin of sRNAs may also lead to cross-mappings, in which a short RNA originating from one locus is partially or completely assigned to a different location (19). Moreover, most available algorithms are dedicated to studying a single sRNA biotype (20), which further restricts the alignment space and can lead to the assignment of reads from different classes to the one being quantified.

Current implementations for sRNA-Seq quantification can be divided in two categories based on their analysis scope: those that quantify only a single sRNA family, such as miRDeep2 (20), and those pursuing to cover the broad sRNA space such as miRge ((21), sRNAbench (22) and ShortStack (23).

We implemented the sMAll rNa dATa analysis pipElinE (MANATEE) for detection and quantification of known and unknown small RNAs. Manatee is not limited to a single sRNA class and achieves highly accurate results, even for elements residing in heavily repeated loci, by making balanced use of existing sRNA annotation and observed read density information during multi-mapper placement. Additionally, it exploits sRNA-Seq reads to detect expressed unannotated genomic loci that could harbor still unknown small RNA products. The user-friendly pipeline of Manatee returns non-coding RNA (ncRNA) expression counts that can be directly utilized downstream, e.g. for differential expression analysis, rendering it easily integrable in larger bioinformatics workflows.

## Materials and methods

### Multimaps analysis

Here we employed available genomic annotation and Uniquely Aligned Reads (UARs) as a means for guiding multi-mapping reads (or “multimaps”). We performed an initial analysis of 14 distinct human sRNA-Seq libraries derived from hepatoblastoma, liver and heart tissue, embryonic stem cells, as well as MCF7 and HepG2 cell-lines, in order to assess the extent of multimaps and UARs in sRNA-Seq datasets (Table S1). Quality-check and pre-processing of all libraries was performed as in Vlachos *et al.* (24). In brief, dataset quality control was performed using FastQC (25). Cutadapt was used for adapter and contaminant removal (26). Reads were mapped against the GRCh38 human reference assembly. UARs and multimaps with up to 50 genomic positions were retained for further analysis. Clusters of UARs were created across the genome for each sample. UARs were considered as reads mapping uniquely to the genome with one allowed mismatch and Bowtie “best strata” (15). Genomic position of UAR includes the information about the mapping chromosome, strand, start and end position. The minimum density of a UAR cluster was set to one read. Non-coding annotation available in Ensembl v85 (27) and miRBase v21 (28) was used to construct a reference for genomic features. Specifically, lincRNA, Mt-rRNA, Mt-tRNA, processed transcript, rRNA, scRNA, snoRNA, snRNA, sRNA and vaultRNA gene types were derived from Ensembl, while miRNA precursor and mature forms were derived from miRBase. A minimum 1nt overlap was required to assign a read to a specific transcript. All transcripts and UARs were extended by 50nt at each end to allow flexibility in the assignment of reads without adding bias.

Fig. 1A presents the average percentage of UARs, multimaps and unaligned reads across the samples. Five examined cases of positioning multimaps were based on reads with 2 to 17 multimapping regions (Fig. 1B). According to the analyzed cases, a multimap may fall into:

1. unannotated regions of UAR clusters (denoted as blue in Fig. 1B)
2. annotated regions lacking UAR clusters (red)
3. annotated regions that also contain UAR clusters (green)
4. unannotated regions that also lack UAR clusters (orange)
5. annotated regions and regions with UAR clusters with no concordance (pink).

**Figure 1:**
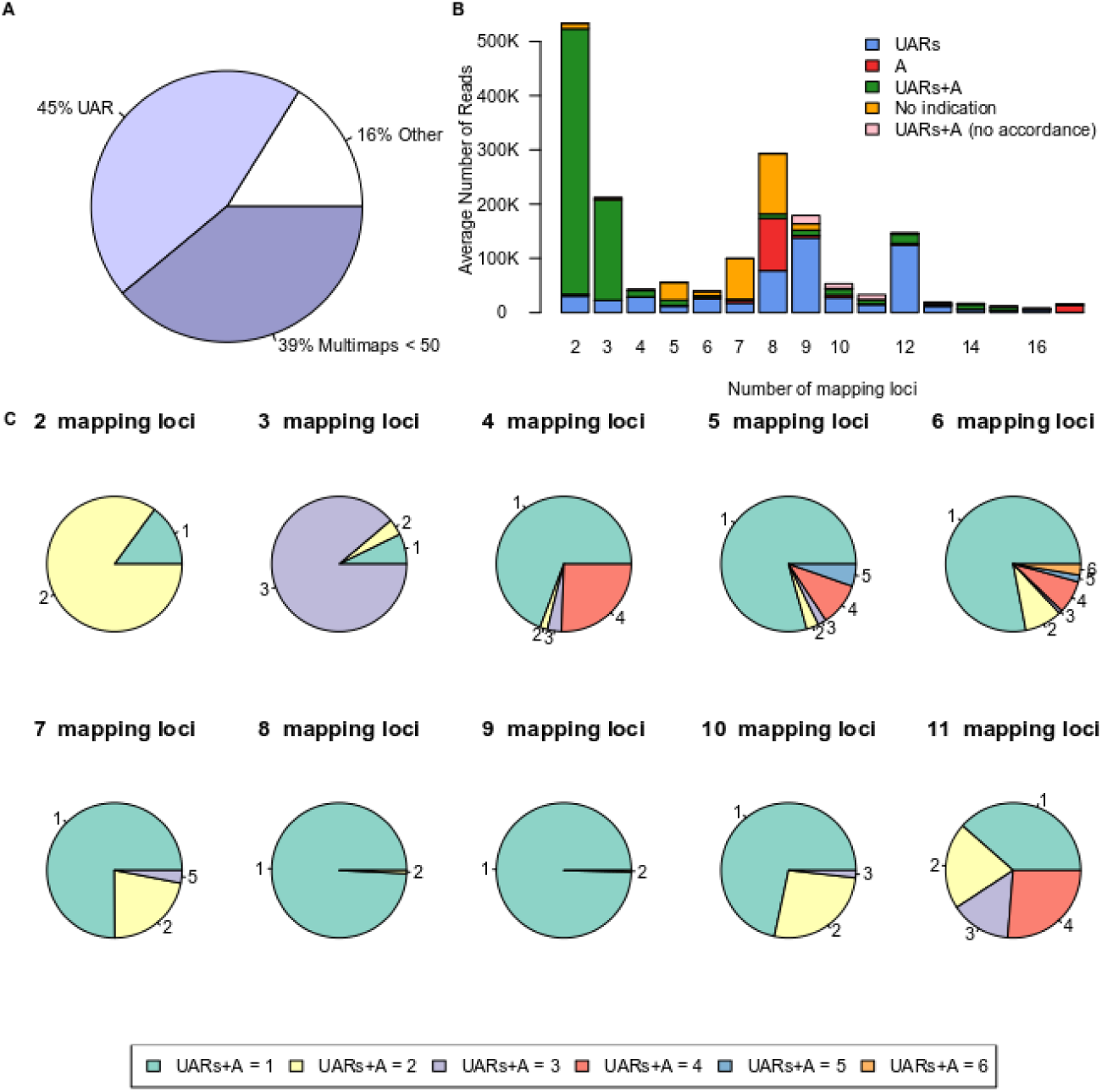
Frequency, proportions and characteristics of multimaps in sRNA-Seq libraries. (A) The average number of UARs, multimaps, and other reads (i.e. unaligned/multimaps exceeding the defined threshold) across all samples. (B) Multimap read categories based on available annotation and UARs. Colors mark five examined cases where each multimap is screened for available annotation and UARs. (C) Proportion of multimaps and the number of their mapping regions with both UAR clusters and available annotation.

Case 3, which includes multimaps falling into regions with both existing annotation and UAR clusters, was further analyzed and examined for the number of such regions per multimap (Fig. 1C). For example, the majority of multimaps with two possible mapping loci had UARs and annotation for both mapping positions. The majority of reads with four possible mapping loci presented one out of four regions with existing UARs and annotation.

A large portion of sRNA-Seq reads (44%) in the analyzed datasets mapped to more than one genomic loci (Fig.1 A). 17.4% of total multimaps fell into regions with UARs lacking annotation and for 13.2% no straightforward information of positioning or annotation was available (Fig. 1B). Algorithms based on genomic alignment which rely entirely on UAR information, may fail to account for cases of multimaps that could otherwise be assigned to existing annotation (red in Fig. 1B, 9.1% of total multimaps). On the other hand, multimaps assigned to more than one genomic features using annotation from a broader spectrum of non-coding RNAs (Fig. 1C) showed that tools dedicated entirely to a specific RNA biotype may be biased towards that type (Fig. S1).

The above conclusions constituted the basis for the Manatee algorithm which attempts to approach the multimap issue by simultaneously incorporating information from UARs and existing annotation.

### Algorithm

#### Input

Manatee requires FASTQ/FASTA sRNA-Seq data files that have been preprocessed for 3’ adapter and barcode removal. Genomic annotation for ncRNAs is required as input in GTF format with the following tags in the attributes field: gene_name, gene_id and gene_biotype.

#### Alignment and quantification

The full outline of abundance estimation adopted by Manatee is provided in Fig. 2. Mapping of sequencing reads is carried out using Bowtie aligner (15). In the primary phase, reads aligned uniquely to the genome are used to form the UAR clusters across the genome. Multimaps are assigned to loci based on the following approach:

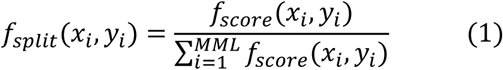

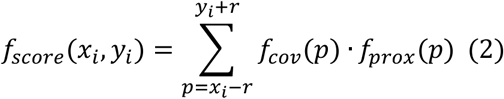

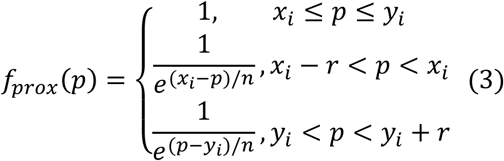

where *x*_*i*_ and *y*_*i*_ are the start and end placement positions of the multimap *i* and *r* is the range in the close proximity of the read (default 50). Function *f*_*cov*_ denotes the UAR density at genomic position *p* and *f*_*prox*_ assigns weights to *f*_*cov*_ based on the position *p* within the genomic region [*x*_*i*_ - r, *y_i_ + r*]. The multimap is split across its valid multi-mapped loci (*MML)* according to the score calculated using function *f*_*split*_. *n* denotes the relevance of approximate density distribution and is set by default to 10. For multimaps with non-matching annotation and positioning of UAR clusters, annotation is preferred and used to guide the final placement of the reads. If a multimap falls into regions which are annotated completely or partially, all relevant transcripts are noted in the output file in the form of alternative transcripts. In case where at least one annotated miRNA is present among those features, the read is assigned to the transcript which exhibits the highest coverage score (ratio):

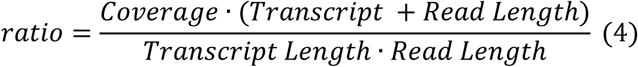

**Figure 2:**
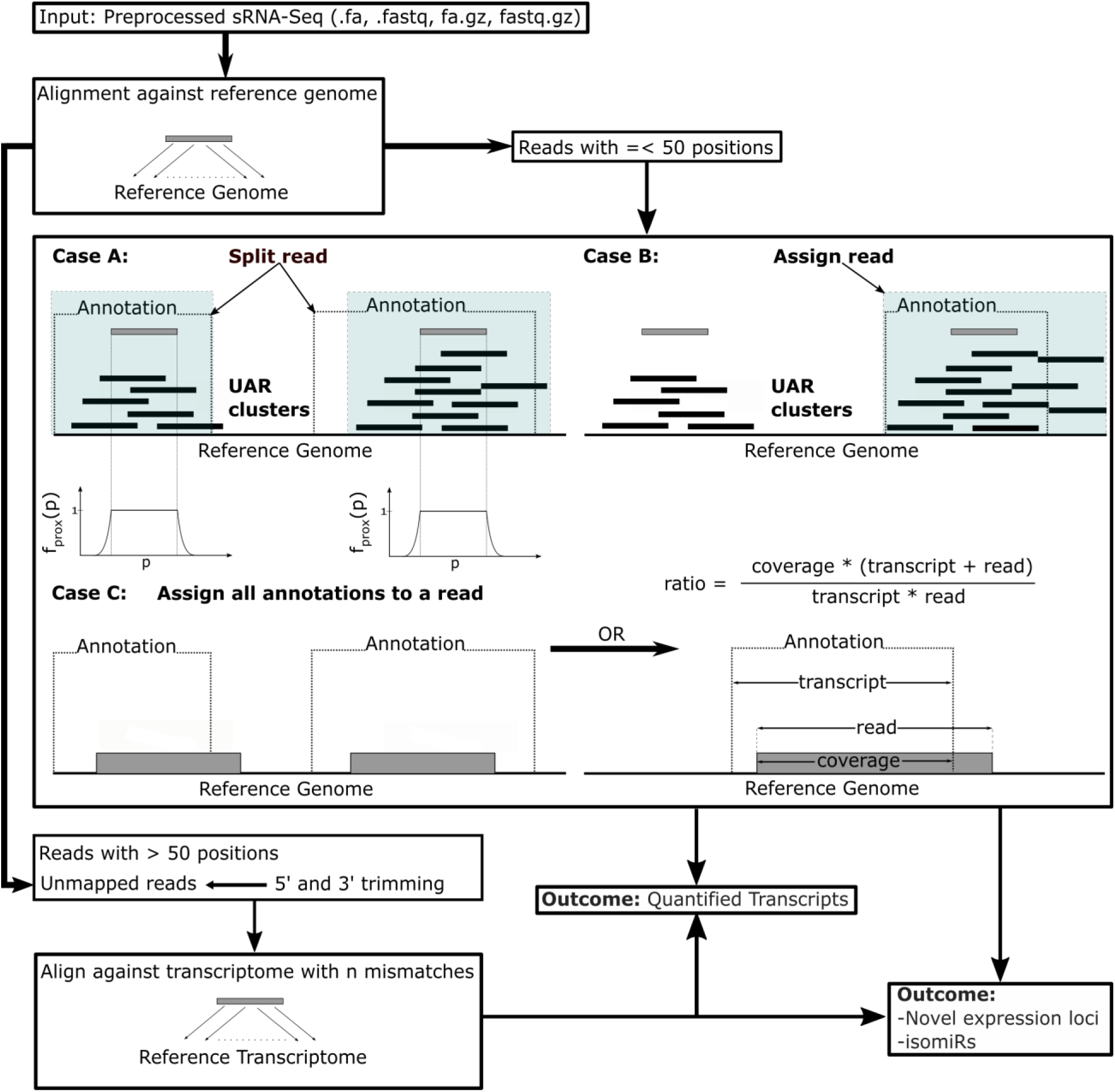
Manatee workflow. Reads with up to 50 multi-mapping positions are either (A) split among their annotated and uniquely aligned read (UAR)-containing loci according to Equation 1, (B) assigned to regions containing both annotation and UARs or (C) assigned to loci with existing annotation. In case of (C), if an annotated miRNA is within the annotated loci, a ratio for selecting the best fitted transcript is used to prioritize mature miRNAs over precursors. Reads with more than 50 mapping positions, reads which could not be mapped to the genome, and reads that could not be assigned to regions with existing annotation and UARs are aligned against the transcriptome with gradual increment of allowed mismatches. The output results contain quantified transcripts, putative novel expression loci and isomiR sequences.

Coverage is the number of overlapping nucleotides between the annotated feature (transcript) and the read length. The ratio heuristic prioritizes the annotation with the highest coverage, while considering read and transcript lengths.

#### Salvaging reads by secondary transcriptome alignment

Reads that exceed the multi-mapping threshold and reads that could not be mapped to the genome are additionally aligned against the transcriptome based on the provided annotation. In the latter case, the number of allowed mismatches is gradually augmented (maximum default 3). In both cases, reads that can be assigned to transcripts with existing mapping densities calculated in previous steps are assigned to those transcripts. If no expression estimates exist, up to five transcripts with the highest mapping quality are retained and assigned as alternative transcripts.

#### IsomiR detection

All reads assigned and quantified as miRNA type are retained and stored in a separate output file. Each detected putative isomiR sequence is stored independently along with its estimated count. Those results may serve in downstream isomiR investigations.

#### Detection of novel unannotated expression loci

UARs mapping to loci lacking genomic features are organized into read clusters based on their genomic positions. Manatee identifies clusters as genomic regions which contain at least five reads and no gap longer than 50nt between consecutive reads with the default parameters, which can be altered by the user. The output of this step is a single file comprising the unannotated genomic loci and their associated read counts.

### Simulated Data

A simulated short read dataset (https://github.com/jehandzlik/Manatee/tree/master/simulated_data) was created using random sampling with a Monte Carlo inversion technique. Human annotation was derived from Ensembl v85 (27), GtRNAdb (29) and miRBase v21 (28). Three randomly selected sRNA-Seq libraries (Table S2) obtained from Gene Expression Omnibus (30) were also employed in the process. Samples were aligned against GRCh38 human reference assembly after 3’-adapter sequences were removed using Cutadapt (26). Since processed sRNA fragments/features are derived from their precursors by biogenesis/cleavage mechanisms that are distinct to each biotype, simulated reads were designed to follow this rationale. Based on uniquely aligned reads observed in the real data, probability mass functions (PMFs) were created for each biotype describing the read start positions. Eight different PMFs were created for the following types: miRNA, tRNA, Mt-tRNA, rRNA, snRNA, snoRNA, lincRNA and non-coding transcript. Likewise, SNPs and read lengths for each sRNA type were also estimated based on PMFs of UARs. More details on the creation process and the dataset characteristics are available in the Supplementary File 1 (Section “Simulated Reads Analysis” and Supplementary Fig. 2-4).

## Results and discussion

### Comparison with other methods using simulated data

The accuracy of Manatee was evaluated using simulated short read dataset (Supplementary Dataset 1). Bowtie v1 (15) was used as a baseline, since it is a commonly used aligner in sRNA-Seq pipelines, while miRge (21), ShortStack (23), and sRNAbench (22) were employed as state-of-the-art approaches in the evaluation. Among many of the available tools for small RNA analysis, miRge performs read alignment entirely against sRNA annotation, ShortStack emphasizes extensively on the genomic alignment of short reads, while sRNAbench (22) extends the functionality of miRanalyzer (29) by applying genomic/transcriptomic alignment of multiple sRNA types in a hierarchical step-wise manner. Those diverse approaches of quantifying sRNAs constitute attractive candidates for direct comparisons with the Manatee algorithm. miRge, ShortStack, sRNAbench (executed using the genomic alignment mode), and Manatee were executed under their default settings. Bowtie was executed permitting a maximum of 1 mismatch and up to 5 multimaps, while transcript quantification was performed with HTSeq-Count (12) using the intersection-nonempty mode and “nonunique all” parameter. The selected parameters for both Bowtie and HTSeq-Count were found to be optimal for the input in question.

Estimated sRNA counts for HTSeq-Count, Manatee, miRge, ShortStack and sRNAbench were contrasted to the ground truth (i.e. simulated counts) (Fig. 3A). All tools tend to over-estimate numerous transcripts that have zero abundance in the simulated dataset (Fig. 3A, Sim.=0 & Est.>=5). However, the opposite behavior was observed at the other end of the spectrum; expressed, and highly expressed transcripts were not assigned any reads (Fig. 3A, Sim.>5 & Est.=0). Among the tested tools, counts estimated by Manatee appeared closest to the simulated abundances (Fig. 3).

**Figure 3:**
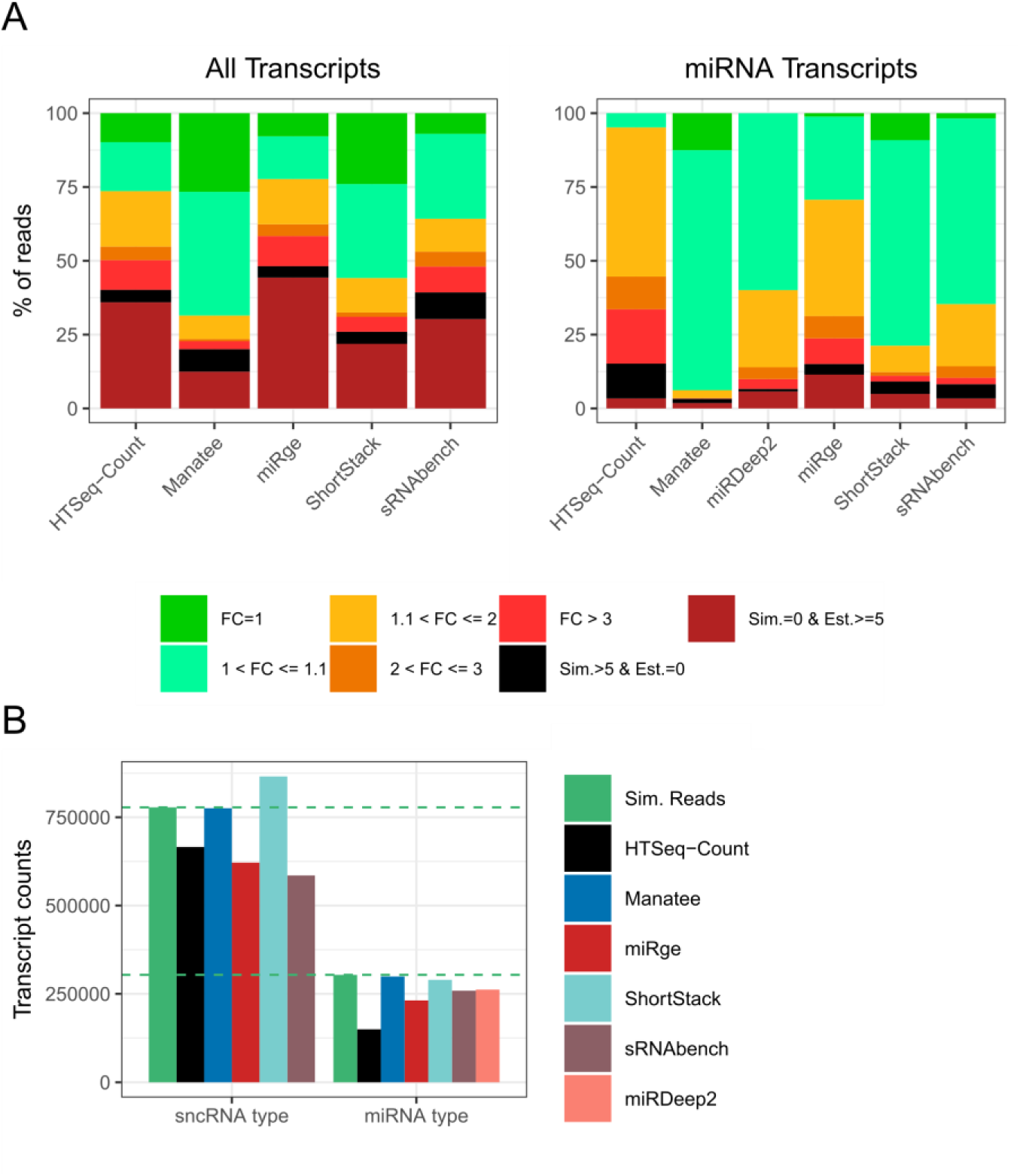
Tools evaluation statistics for simulated sRNA-Seq data. (A) Fold changes for simulated vs. estimated transcript counts for evaluated tools utilizing all small ncRNA species or only miRNAs. Fold change of 1 denotes no difference between the simulated and the calculated counts. Sim. > 5 & Est.=0 denotes percentage of reads where the simulated transcript counts > 5 were estimated as zeros by the examined tools. Sim.=0 & Est. > =5 relates with proportion of estimated transcript counts > 5 for which the true simulated count was zero. (B) Comparison between the ground truth count sum of simulated reads and the total estimated transcript counts across implementations.

Manatee is not only able to map and accurately quantify diverse sRNA classes, but fares favorably when compared to methods specifically designed for miRNAs. miRDeep2 (20), which uses Bowtie to map sequencing reads against precursors and discards or assigns multimaps equally to their valid loci, was executed against the same dataset, with default settings. These results vividly depict that Manatee users can quantify and investigate still underexplored small RNA classes, while having accurate and robust results for miRNAs, which is the super star small ncRNA class. Sum of simulated transcript counts was contrasted against the estimated counts by the six tools. ShortStack displayed tendency for count inflation, while miRge, miRDeep2 and sRNAbench underestimated transcript counts (Fig. 3B). Precision metrics for the examined algorithms were estimated and are presented in Table 1 and Figure S5.

**Table 1.** Performance metrics for evaluated implementations.

| Table 1. Performance metrics for evaluated implementations |  |  |  |  |  |  |
| --- | --- | --- | --- | --- | --- | --- |
| Tool | RNA type | RMSD | Jaccard distance | Euclidean distance | Pearson correlation | Spearman correlation |
| HTSeq-Count | small ncRNAs | 372.614 | 0.298 | 20439.478 | 0.798 | 0.577 |
| Manatee |  | **341.494** | **0.173** | **15730.981** | **0.879** | **0.796** |
| miRge |  | 408.08 | 0.503 | 26553.417 | 0.641 | 0.581 |
| ShortStack |  | 456.499 | 0.271 | 20939.313 | 0.801 | 0.655 |
| sRNAbench |  | 361.399 | 0.395 | 22805.388 | 0.744 | 0.556 |
| HTSeq-Count | miRNAs | 369.164 | 0.442 | 8813.660 | 0.529 | 0.392 |
| Manatee |  | **107** | **0.031** | **2556.831** | **0.929** | **0.954** |
| miRge |  | 249.67 | 0.216 | 6419.008 | 0.731 | 0.684 |
| ShortStack |  | 236.657 | 0.151 | 5738.626 | 0.683 | 0.737 |
| sRNAbench |  | 215.574 | 0.138 | 5218.504 | 0.752 | 0.725 |
| miRDeep2 |  | 155.313 | 0.078 | 3867.274 | 0.893 | 0.827 |

### Comparison with other methods using real sRNA-Seq data

sRNA-Seq data derived from breast cancer MCF7 cells (Study ID: SRP060224, Sample ID: SRR2084358) was utilized to cross-correlate the compared sRNA/miRNA quantification methods (Fig. 5) using Pearson correlation. Seven genomic features exhibiting read counts above 10,000 reads for all executions were removed from the comparison as outliers (Table S3).

**Figure 5:**
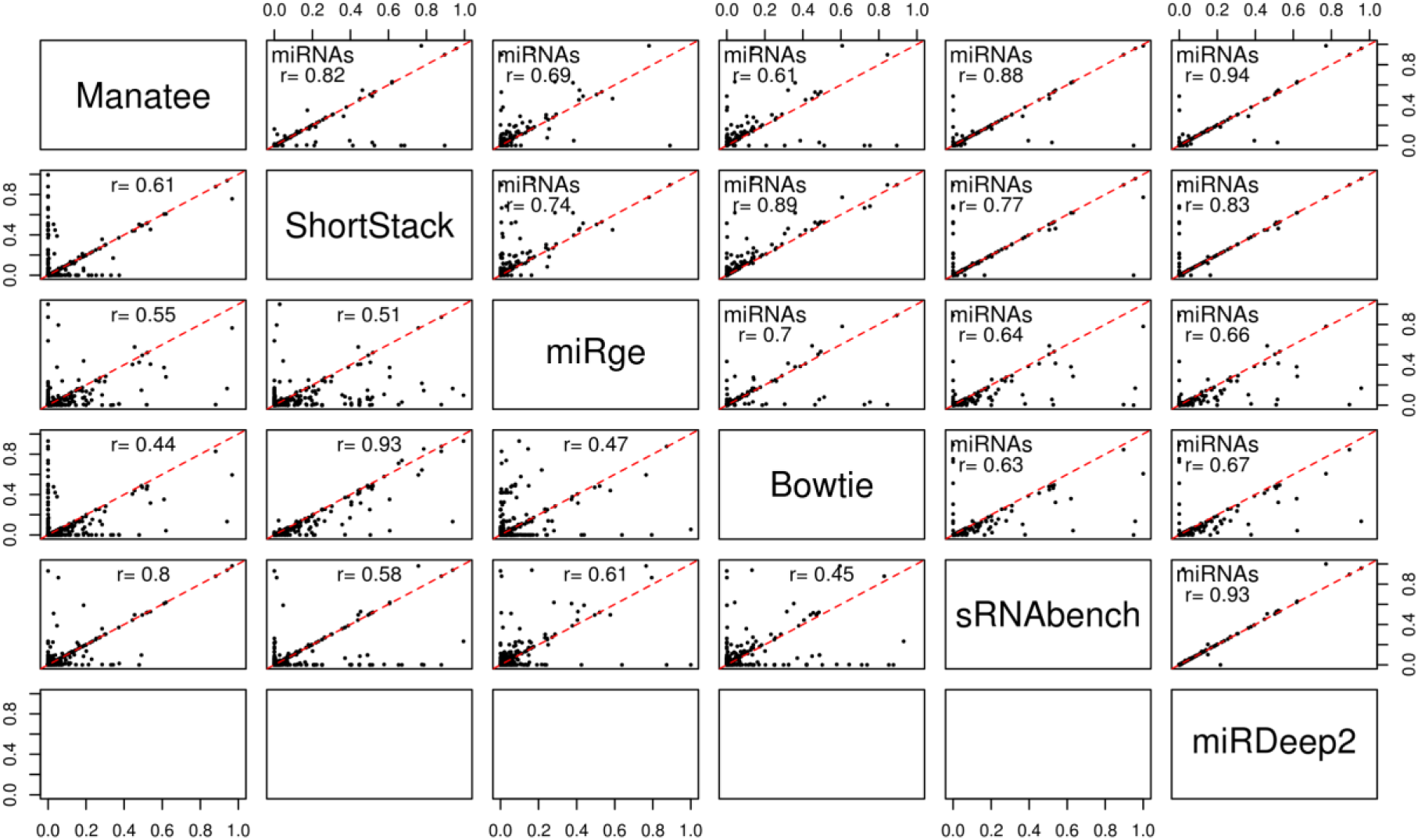
The analyzed sRNA-Seq sample was compared across 5 methods for all sRNA types (lower left panel) and across 6 methods for miRNAs (upper right panel). Pearson correlation was calculated for each pair of compared tools and denoted on each plot with the red line indicating the perfect correlation.

This comparison enabled the assessment of concordance between tools when using real data. For miRNAs, Manatee exhibited > 0.8 correlation coefficient with ShortStack, sRNAbench and miRDeep2, whereas miRge and Bowtie+HTSeq-Count displayed similar patterns of underestimating transcript abundance, as occurred in comparisons using simulated data. ShortStack results also appeared on par with the findings of the previous section, where a high number of false positive and negative counts was present. When comparing the total sRNA transcriptome results, a substantial divergence between the estimated counts was observed across executions. These findings may indicate the existence of intrinsic properties in each tool that, in some cases, drive to misclassification and erroneous quantification of sRNAs.

### Unannotated clusters

Manatee supports the detection of expressed unannotated loci that can be used to identify novel sRNAs and sRNA classes in diverse research settings. Execution of Manatee with default settings on the MCF7 sRNA-Seq sample (Study ID: SRP060224, Sample ID: SRR2084358) detected a total of 588 unannotated clusters. 503 clusters with cluster length < 50nt are shown in Figure 6 (mean reads per cluster µ=35.06 and σ^2^=114.05). Users aiming to proceed with the detection of novel sRNA genomic loci, are strongly advised to first overlap the detected clusters with protein coding exon annotation in order to exclude putative products of mRNA degradation events (30). Following this filtering step and using coding annotation derived from Ensembl v85, 74 clusters remained as highly promising loci for further investigation.

**Figure 6:**
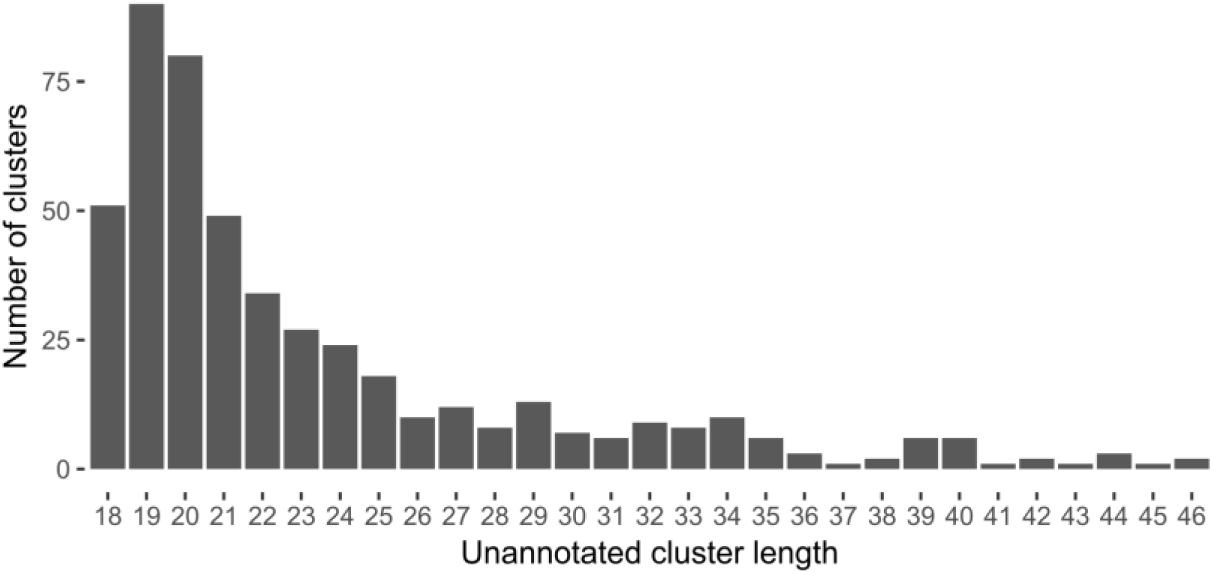
Length distribution of uniquely aligned read clusters lacking annotation in real sRNA-Seq sample.

## Conclusion

Small RNA-Seq experimental datasets require extra caution during the alignment and quantification processes compared to RNA-Seq libraries due to technical obstacles arising from small read and transcript lengths. Short sequences tend to map to more than one genomic region, thus affecting transcript quantification. Variations and post-transcriptional modifications introduce an additional layer of complexity to the detection of proper mapping loci. Many available tools for sRNA-Seq analysis focus on specific short RNA types, at the expense of other subclasses and the breadth of the investigation. Alignments solely against transcripts neglect expression patterns which could bear information about their origin. Manatee adopts a step-wise approach, exploiting (i) available annotation and (ii) reliable/robust density information towards an optimized multi-mapping read placement. Importantly, it enables the detection and quantification of putative expressed small RNA loci lacking annotation. Future expansions of the algorithm could include the incorporation of tolerance against common post-transcriptional modifications or indels to further boost the precision of transcript quantification in a broader and more realistic alignment space.

Manatee provides an improved approach to quantify transcripts present in sRNA-Seq data by combining reliable information inferred from UARs and transcript annotation, to more accurately guide the placement of multi-mapping reads. It is an efficient and user-friendly tool that can be a significant aid in small RNA studies.

## Data availability

Project name: Manatee 1.0 Project home page: https://github.com/jehandzlik/Manatee.

## Acknowledgements

We thank Artemis Fakorelli for editing the manuscript.

## Author information

### Contributions

J.E.H designed the algorithm and performed further analysis. S.T. tested the algorithm and executed tool comparison. I.S.V. and A.G.H. supervised the research. J.E.H, S.T. and I.S.V. wrote the manuscript. All authors reviewed and approved the manuscript.

### Corresponding author

Correspondence to Artemis Georgia Hatzigeorgiou.

## Supplementary information

Supplementary file 1, Supplementary Dataset 1

