## Supplementary File for "Manatee: detection and quantification of small non-coding RNAs from next-generation sequencing data"

[1]DIANA-Lab, Department of Electrical & Computer Engineering, University of Thessaly, Volos 38221, Greece, [2]Department of Biology, University of North Dakota, Grand Forks, North Dakota 58202, U.S.A., [3]Hellenic Pasteur Institute, Athens 11521, Greece, [4]Harvard Medical School Initiative for RNA Medicine, Department of Pathology, Cancer Research Institute, Beth Israel Deaconess Medical Center, Harvard Medical School, Boston, Massachusetts 02115, U.S.A.

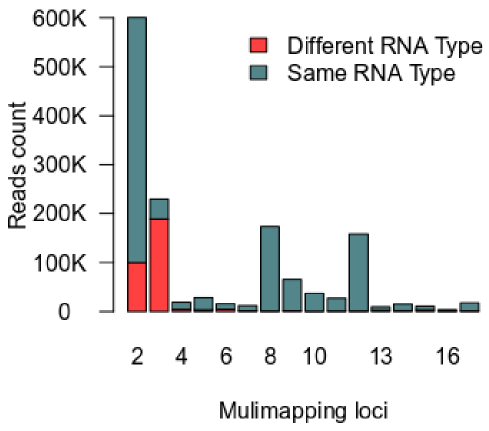


*Supplementary Fig. 1: Concordance between multimapping loci and corresponding biotypes. Proportion of multimapping reads with associated annotation of the same or different RNA biotype for each of the multimapping loci. A substantial number of reads with two and three multimapping positions associate with annotations of dissimilar small RNA types.*

#### Simulated Reads Analysis

Simulation of sRNA-Seq reads was based on parameters and statistics estimated from three randomly selected sRNA-Seq libraries (Table S2). The proportion of sRNA biotypes present in the simulated dataset was based on predefined probabilities (Fig. S2A). The start position of reads for each genomic feature was sampled from cumulative distribution functions (CDFs) created for each sRNA biotype separately and based on the positioning of UARs in the dataset utilized in the simulation (Fig. S3). To determine the uniquely aligned clusters of reads, the input data was pre-processed as in Vlachos *et al.* (1) and aligned against the GRCh38 genome assembly using Bowtie v1 (2) with 1 allowed mismatch. Reads presenting ≥20% overlap with a specific genomic feature were assigned to that feature and used in the CDF estimation. For example, according to the generated miRNA CDF, simulated reads for this biotype have a high probability (87%) to start at position 0 of the mature miRNA annotation. The probability of a simulated read to bear a sequence modification was based on the observed 3' and 5' modifications (non-templated additions) and SNPs in UARs in the examined sRNA-Seq libraries (Fig. S2B-C). Similarly to the case of the read starting positions, SNPs, non-template additions and read lengths were also sampled from generated CDFs based on the respective information retrieved from UARs (Fig. S4). Figure S4A-B present the CDFs of SNP positions and transcript lengths excluding the lengths of miRNA type, while panels 4C-D display the probabilities of non-template modification types. Lengths of miRNA transcripts were sampled from a truncated normal distribution with m=22, s=1, a=18 and b=24. Read counts generated for each transcript were sampled from negative binomial distribution corresponding to selected for simulation transcript biotype. Count of the simulated read $X_{ij}$ for transcript *i* and biotype *j* was sampled from

$$X_{ij} \sim Negative\ Binomial\ (mean = \mu_{ij}, variance = \mu_{ij} + \frac{1}{r_{ij}}\mu_{ij}^2)$$

where $\frac{1}{r_{ij}}$ is a dispersion parameter which was set to $r_{ij} = \frac{\mu_{ij}}{4}$ by default for every RNA biotype used in the simulation. Mean $\mu_{ij}$ of transcript *i* and biotype *j* was estimated based on the UARs present in the input libraries which mapped to selected for simulation biotypes.

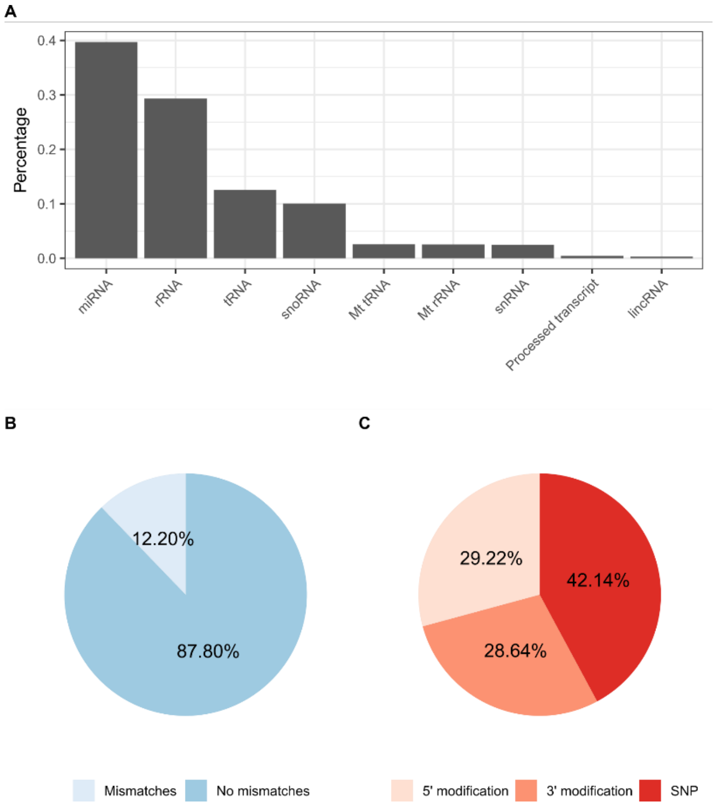


*Supplementary Fig. 2: sRNA biotype distribution present in the simulated data and statistics of sequence modifications in the selected sRNA-Seq libraries. (A) Proportion of sRNA biotypes present in the simulated dataset. (B) Proportion of reads with mismatches and (C)SNPs, 3' and 5' non-templated additions observed across the sRNA-Seq libraries used in the simulation process.*

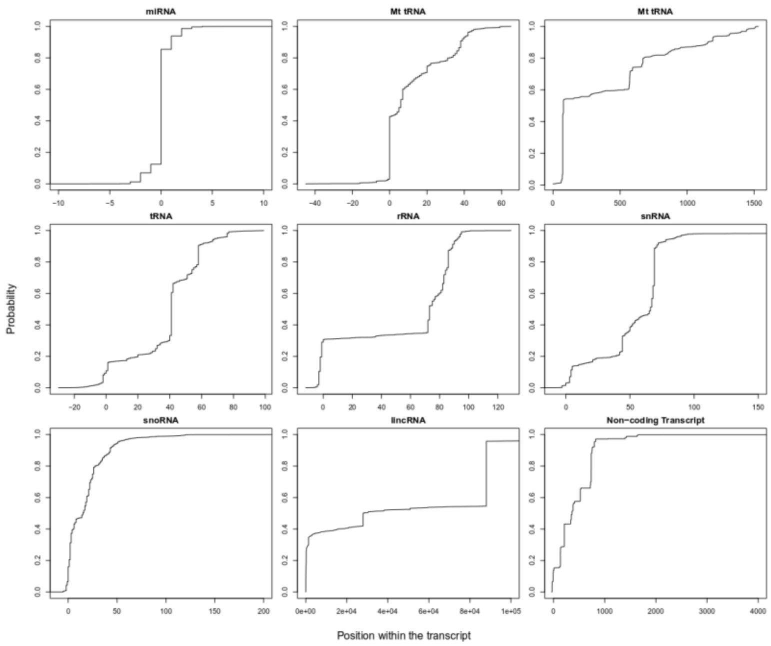


*Supplementary Fig. 3: CDFs of starting position of reads per biotype, based on UARs present in the sRNA-Seq libraries selected for simulation.*

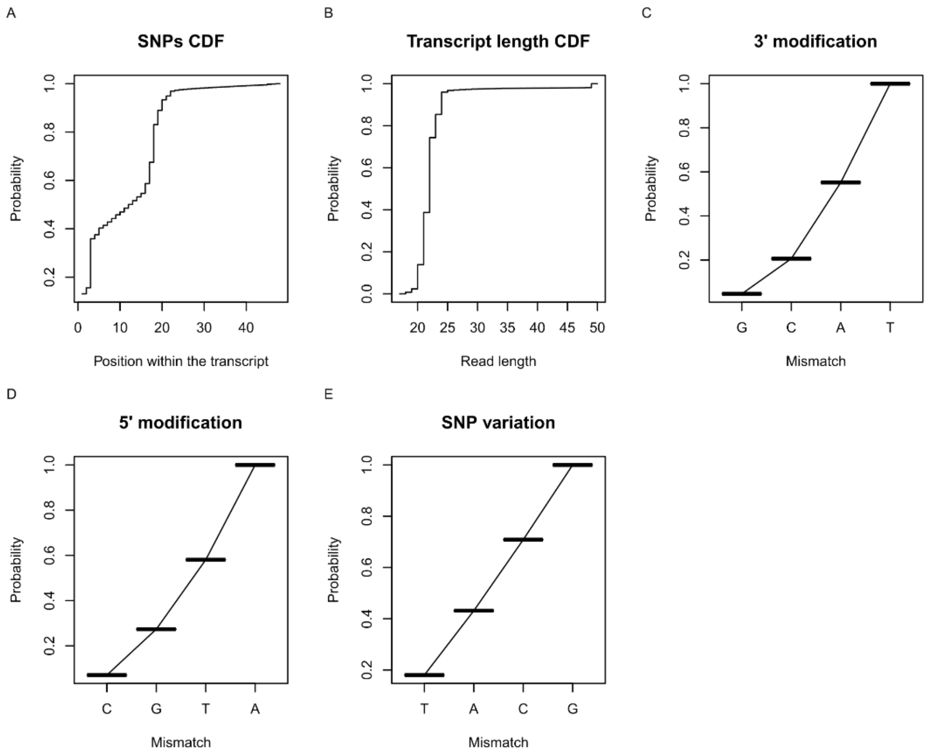


*Supplementary Fig. 4: CDFs of SNP positions, read lengths and probabilities of SNPs, non-template 3' and 5' additions, as estimated from the UARs present in the selected sRNA-Seq libraries. (A) Estimated CDF for the SNP position within the transcript area. (B) Estimated CDF for read length excluding miRNA lengths. Estimated probabilities for (C) 3' or (D) 5' non-template additions and for SNP nucleotide variation (E).*

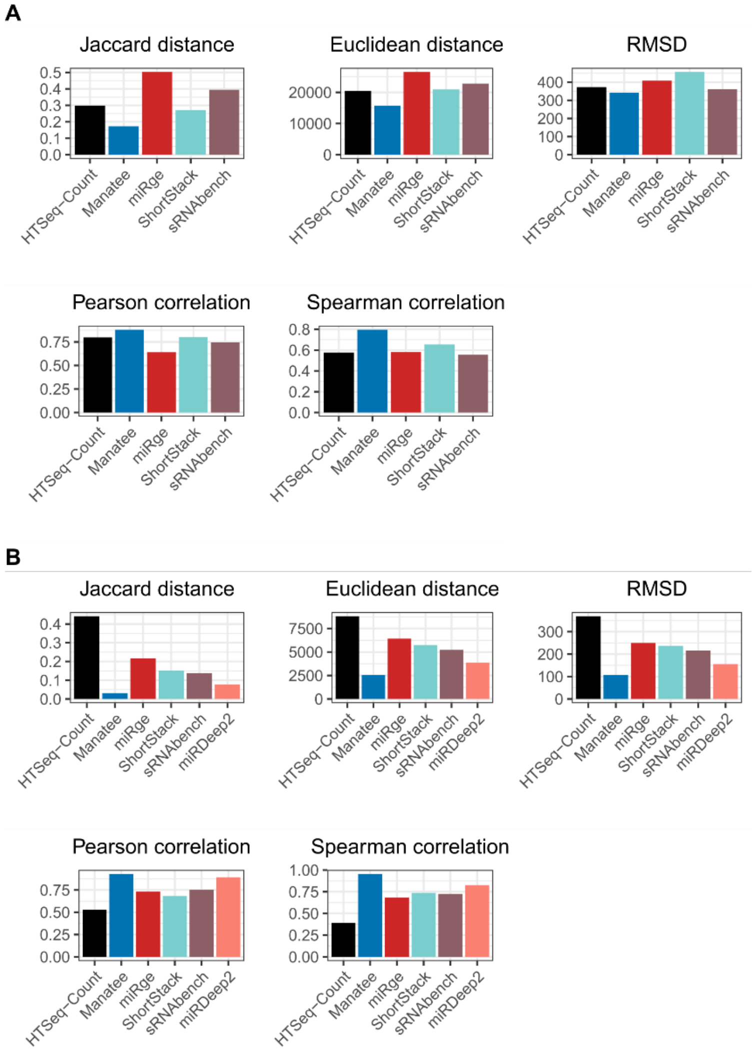


*Supplementary Fig. 5: Precision metrics estimated using the simulated dataset: Jaccard distance, Euclidean distance, Root-mean-square deviation (RMSD), Pearson and Spearman correlation for counts estimated by compared tools for (A) small ncRNAs and (B) miRNAs.*

### Supplementary Tables

#### Table S1: sRNA-Seq datasets utilized in the multimaps analysis along with the related study and sample ID, and the relevant metadata information (cell/tissue type, sample type, and condition).

| **Supplementary Table S1.** Real sRNA-Seq datasets utilized in the multimap analysis | | | | | |
|---|---|---|---|---|---|
| **#** | **Study ID** | **Sample ID** | **Cell/Tissue Type** | **Sample Type** | **Condition** |
| **1** | SRP049449 | SRR1636959 | hESC | Cells | Healthy |
| **2** | SRP049449 | SRR1636960 | hESC | Cells | Healthy |
| **3** | SRP060224 | SRR2084358 | MCF7 | Cell-lines | Breast cancer |
| **4** | SRP060224 | SRR2084359 | MCF7 | Cell-lines | Breast cancer, DHA-treated |
| **5** | SRP065616 | SRR2889755 | HepG2 | Cell-lines | Liver cancer |
| **6** | SRP065616 | SRR2889756 | HepG2 | Cell-lines | Liver cancer |
| **7** | SRP049590 | SRR1642941 | Liver | Tissue | Healthy |
| **8** | SRP049590 | SRR1642943 | Liver | Tissue | Healthy |
| **9** | SRP058632 | SRR2038050 | Liver | Tissue | Healthy |
| **10** | SRP058632 | SRR2038051 | Liver | Tissue | Healthy |
| **11** | SRP048606 | SRR1596226 | Liver | Tissue | Hepatoblastoma |

| | | | | | |
|---|---|---|---|---|---|
| **12** | SRP048606 | SRR1596227 | Liver | Tissue | Hepatoblastoma |
| **13** | SRP058632 | SRR2038044 | Heart | Tissue | Healthy |
| **14** | SRP058632 | SRR2038045 | Heart | Tissue | Healthy |

**Table S2: sRNA-Seq datasets utilized in the simulation process along with the related study and sample ID, and the relevant metadata information (cell/tissue type, sample type, and condition).**

| **Supplementary Table S2.** sRNA-Seq datasets utilized in the simulation process. | | | | | |
|---|---|---|---|---|---|
| **#** | **Study ID** | **Sample ID** | **Cell/Tissue Type** | **Sample Type** | **Condition** |
| **1** | SRP059434 | SRR2061810 | Liver | Tissue | Healthy |
| **2** | ERR842903 | ERR842903 | Gallbladder | Tissue | Healthy |
| **3** | SRP006574 | SRR191548 | Breast | Tissue | Healthy |

**Table S3: Transcripts excluded from the comparison analysis for real sRNA-Seq data. Seven transcripts with read count > 10,000 estimated by each examined tool were considered as outliers and removed from the comparison.**

| **Supplementary Table S5.** Transcripts excluded from the comparison analysis for real sRNA-Seq data | | | | | | |
|---|---|---|---|---|---|---|
| **Transcript** | **Bowtie** | **miRge** | **ShortStack** | **Manatee** | **sRNAbench** | **miRDeep2** |
| hsa-miR-26a-5p | 12,826 | 13,004 | 12,968 | 13,020.895 | 12,980 | 12,961 |
| hsa-miR-21-5p | 76,203 | 229,645 | 229,014 | 229,881.081 | 229,372 | 228,759 |
| hsa-miR-125a-5p | 14,408 | 13,212 | 14,587 | 14,654.986 | 14,622 | 14,573 |
| hsa-miR-99b-5p | 43,811 | 43,200 | 47,015 | 47,224.991 | 47,109 | 46,969 |
| hsa-miR-191-5p | 32,305 | 33,359 | 34,394 | 34,601.992 | 34,494 | 34,351 |
| hsa-miR-182-5p | 20,083 | 20,188 | 26,060 | 26,152.996 | 26,044 | 26,010 |
| hsa-miR-27b-3p | 77,089 | 77,385 | 79,150 | 79,562.239 | 79,078 | 78,963 |
